# FANCY: Fast Estimation of Privacy Risk in Functional Genomics Data

**DOI:** 10.1101/775338

**Authors:** Gamze Gürsoy, Charlotte M. Brannon, Fabio C.P. Navarro, Mark Gerstein

**Affiliations:** Program in Computational Biology and Bioinformatics, Yale University, New Haven, CT 06520, USA; Department of Molecular Biophysics and Biochemistry, Yale University, New Haven, CT 06520, USA; Department of Computer Science, Yale University, New Haven, CT 06520, USA

## Abstract

Functional genomics data is becoming clinically actionable, raising privacy concerns. However, quantifying the privacy leakage by genotyping is difficult due to the heterogeneous nature of sequencing techniques. Thus, we present FANCY, a tool that rapidly estimates number of leaking variants from raw RNA-Seq, ATAC-Seq and ChIP-Seq reads, without explicit genotyping. FANCY employs supervised regression using overall sequencing statistics as features and provides an estimate of the overall privacy risk before data release. FANCY can predict the cumulative number of leaking SNVs with a 0.95 average *R*^2^ for all independent test sets. We acknowledged the importance of accurate prediction even when the number of leaked variants is low, so we developed a special version of model, which can make predictions with higher accuracy for only a few leaking variants. A python and MATLAB implementation of FANCY, as well as custom scripts to generate the features can be found at https://github.com/gersteinlab/FANCY. We also provide jupyter notebooks so that users can optimize the parameters in the regression model based on their own data. An easy-to-use webserver that takes inputs and displays results can be found at fancy.gersteinlab.org.

## Introduction

With the surge of genomics data and the decreasing cost of sequencing [1], genome privacy is an increasingly important area of study. Traditional DNA sequencing, functional genomics [2], and molecular phenotype [3,4] datasets create quasi-identifiers, which, in turn, can be used to re-identify or characterize individuals without their consent. The surge in widely available functional genomics data increases correlations between phenotype and genotype datasets, which amplifies the possibility of re-identification and characterization of the individuals who participate in these studies. Functional genomics data allows for a detailed characterization of disease states and susceptibility, and broad dissemination of this data can promote key scientific advances. Unlike DNA sequencing, functional genomics experiments are not performed for genotyping purposes (rather, for understanding phenotypes and basic biology). Yet, they still yield raw reads containing a substantial amount of patients’ variants, which raises privacy concerns. Thus, there is a trade-off between utility and privacy when it comes to sharing functional genomics data. This trade-off can be difficult for scientists to navigate. For example, scientific funding agencies’ data-sharing and privacy policies about controlled-access can prevent the release of data, sometimes as late as after the relevant article has published due to concerns about privacy [5].

In contrast to DNA sequencing-based data such as GWAS, few tools currently exist to assess the risk of privacy loss from functional genomics data. In order to protect patient privacy while promoting scientific progress through data-sharing, it is essential that we develop robust methods for making these assessments. Such assessments may bolster informed consent and empower scientists to plan functional genomics experiments with patient privacy in mind.

Before the release of data from a functional genomics experiment, it is essential to be able to rapidly quantify the number of leaking variants. This is particularly important when multiple assays are performed on samples from the same individuals. That is, a single data type may not leak enough variants for privacy to be a concern, but a combination of different functional genomics data can pose significant privacy risk, as different assays target different regions of the genome with different coverage profiles [e.g., RNA sequencing (RNA-Seq) targets expressed exons, whereas H3K27ac chromatin immunoprecipitation sequencing (ChIP-Seq) targets the non-coding genome on the promoter and enhancer regions] and depth profiles (i.e., some assays have spread out peaks while others are more punctuated). Such quantification is possible by genotyping the raw sequences and overlapping them with the gold-standard genotypes (e.g. those obtained from whole-genome sequencing) from the individuals. The use of a gold-standard is necessary because functional genomics data alone provides a less reliable picture of an individual’s genotypes, and may lead to false positives due to the targeted nature of the assays. The limitations of this approach are (i) the large resources required for genotyping; in principle it is possible to genotype the raw reads with current genotyping tools, but an average variant calling pipeline would need to be radically re-parameterized to suit different assays, as traditional genotyping softwares are typically optimized for whole-genome sequencing and (ii) the need for a gold-standard genotype data belonging to the patient, which may not be readily available in every case.

In this study, we developed FANCY, a supervised learning method to infer the number of leaking single nucleotide variants (SNVs) from raw functional genomics data. Our primary goal was to quantify the SNV leaks in raw sequences without needing genotyping or a gold-standard genotype list. We built a Gaussian Process Regression model that takes the assay type, sequencing features such as depth 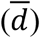 and breadth (*b*) of the coverage, and the statistical properties of the depth distribution such as standard deviation (*σ*), skewness (±*s*), and kurtosis (*k*) as input, and predicts the cumulative number of leaking SNVs. FANCY can separately estimate the number of rare and common variants, and outputs each estimated number with a predicted upper and lower bound in the 95% confidence level. FANCY also outputs a simple qualitative warning message for the risk of sharing (green: can be shared, yellow: attention needed, red: cannot be shared), making it accessible to a range of users. In addition to estimating privacy risk, FANCY can be used to plan functional genomics experiments; for a target number of SNVs, one can back-calculate the required sequencing statistics. The privacy risk assessment is of critical value when the number of leaking variants are low, as in critical ranges, leakage of a few additional variants can change the status of the privacy risk from “can be shared’’ to “cannot be shared’’. Therefore, we also trained our model with data that has SNV leakage fewer than 1000 variants to obtain more accurate results in the lower ranges. This model is called FANCY_low_. A user can first predict the leakage with FANCY. If the number of predicted SNVs is below 1,000, the user can then use FANCY_low_ to fine-tune the accuracy of the prediction.

In addition to FANCY, we also developed a Random Forest classifier plug-in that predicts the type of assay (RNA-Seq vs. ATAC-Seq vs. ChIP-Seq) used to obtain a given dataset by using the sequencing statistics as features. This kind of reverse identification of data may be useful to the community for samples with missing metadata.

## Materials and Methods

### 1. FANCY Details

FANCY is a two-step method. The first step is a regression framework that uses a Gaussian Process Regression (GPR) model with a Matern kernel [12]. We obtained the features as follows: We first aligned the raw functional genomics reads to the reference genome [13] is used for ChIP and ATAC-seq data; STAR [14] is used for RNA-Seq data). We then calculated the depth per base pair using samtools [15] and calculated the following statistics: depth 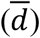 and breadth (*b*) of the coverage, and the statistical properties of the depth distribution such as standard deviation (*σ*), skewness (±*s*), and kurtosis (*k*) (Figure 1a). For the true number of SNVs, we used GATK [8,9] with appropriate parameterization for each assay type) to call the SNVs. After filtering low-quality SNVs as suggested by the Best Practices, we overlapped the remaining SNVs with the gold-standard SNVs generated from whole genome sequencing data to obtain the true number of leaking SNVs (Figure 1c). The second step is the estimation of rare vs. common variants. We divided the 1,000 Genomes data into rare and common categories based on minor allele frequency. For each individual in the 1,000 Genomes Project, given an assay, we found the rare variant density and used the mean density of all individuals with the predicted number of total leaking variants to estimate the number of rare vs. common variants.

**Fig. 1.**
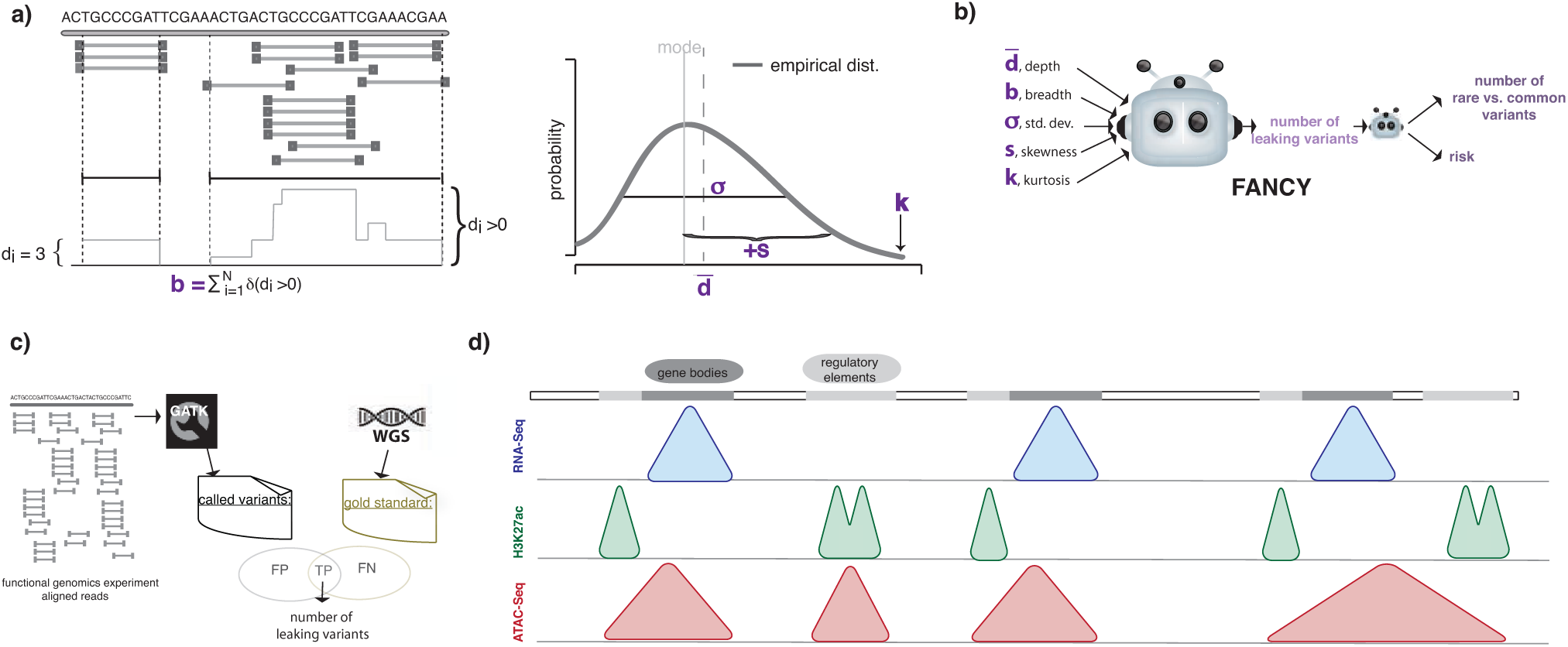
Details of FANCY. (a) The schematic of the features of FANCY: average depth; breadth, i.e number of nucleotides represented with at least one read; first, second and third moment of the depth distribution (standard deviation, skewness and kurtosis). If the distribution is skewed to the right-hand side of the mode (mean is larger than the mode), the skewness is positive. It is negative if the mean is smaller than the mode. (b) The schematic of inputs and outputs of FANCY. (c) The process of determining true number of leaking variants from functional genomics reads. (d) The regions represented by each assay type. The reads of RNA-Seq are concentrated on the gene bodies, H3K27ac ChIP-Seq is concentrated on the non-coding genome (enhancers and promoters), and because ATAC-Seq covers the open chromatin, the reads are concentrated on both coding and non-coding regions.

FANCY_low_ uses the same regressor with the same set of features. However, the training data used in FANCY_low_ contains leaking SNVs up to 1,000 to increase the accuracy when the number of leaking variants is low.

### 2. Gaussian Process Regression

GPR is a supervised learning method that is based on learning fitting functions to a given set of training data. In comparison, traditional regression models learn the parameters of a given function. GPR is a nonparametric method used to calculate the probability distribution over all functions that fit the data instead of calculating the probability distribution of a specific function’s parameters. The advantage of using GPR is its ability to provide uncertainty estimations at a given confidence level. The disadvantage of this method is the computational complexity, which makes it unfeasible for large datasets. Since the sequencing statistics relate to the number of inferred variants differently in different regimes and for different assays (Figure 1c), the relationship between features and the number of leaking variants cannot be modeled by general mathematical approaches such as generalized linear models (Figure 2). A Gaussian process can be defined by its mean and covariance functions as

**Fig. 2.**
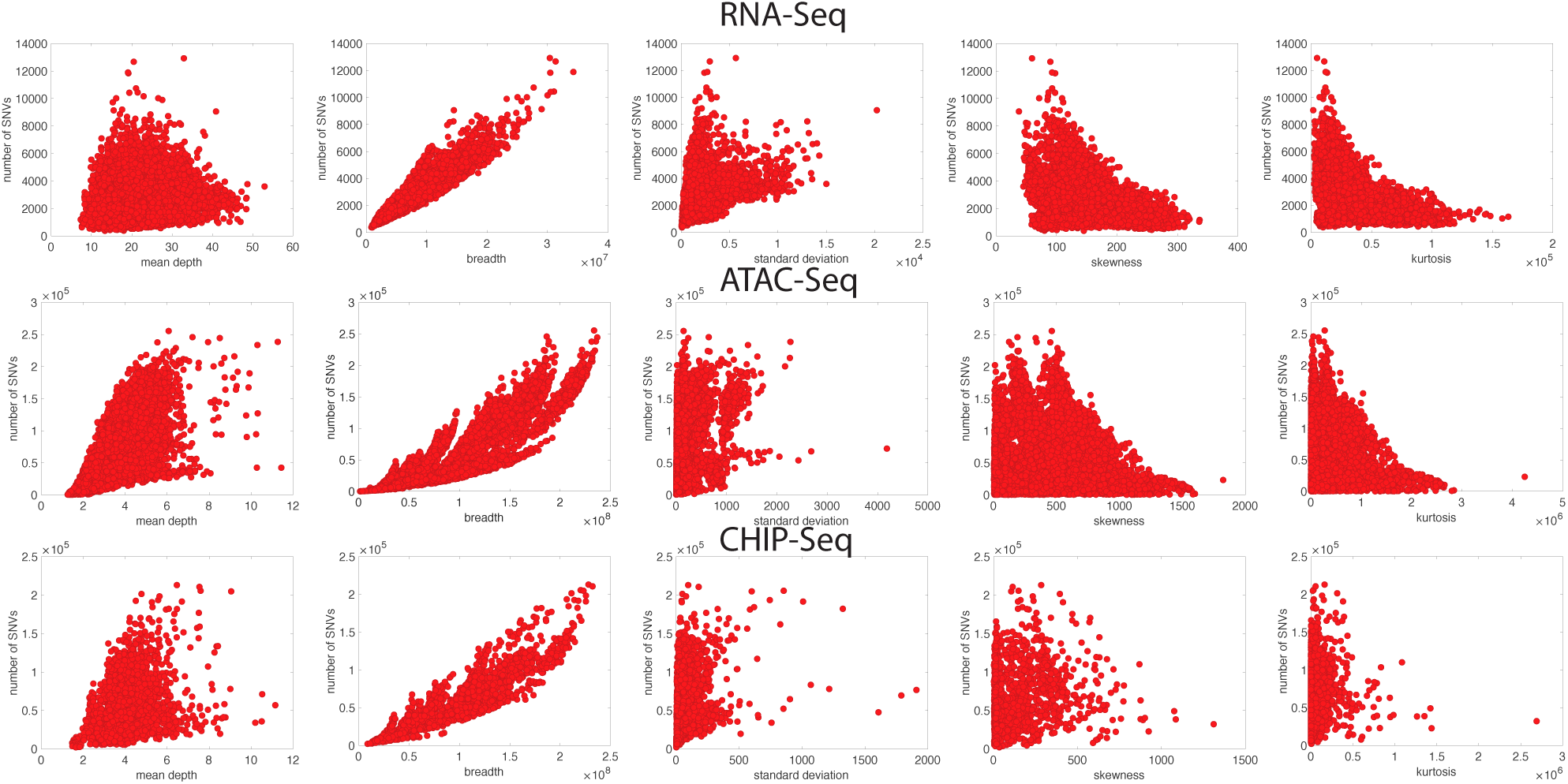
Relationship between sequencing statistics and number of leaking variants. Overall, breadth of the coverage has the highest correlation with the number of leaking variants, while other statistics still show a decent correlation.

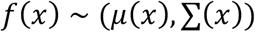

A Gaussian process assumes that the distribution of the values of functions *p*(*f*(*x*_1_), *f*(*x*_2_), = …, *f*(*x*_*N*_)) at a set of points (*x*_1_, *x*_2_, …, *x*_*N*_) is jointly Gaussian with a mean *μ*(*x*) and covariance ∑(*x*), where ∑_*ij*_ = *k*(*x*_*i*_, *x*_*j*_). *K* is a kernel function, which determines the similarity between data points *x*_*i*_ and *x*_*j*_. If these points are deemed similar by the kernel, we expect the output of the functions at these points to be similar as well. For each *x*_*i*_, *x*_*j*_ in our training dataset, we can write a function *f*(*x*_*i*_) such that

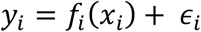

where *ϵ*_*i*_ ∼ *N*(0, *σ*^2^). Therefore, for any input vector (*x*_1_, *x*_2_, …, *x*_*N*_), *f*(*x*) has a joint Gaussian distribution. The covariance (kernel) function *k* is generally taken as Gaussian (i.e., squared exponential kernel). However, in this application, we found that a Matern kernel performs better. We used five-fold cross-validation to avoid overfitting and a separate test dataset to validate our model. We tried other regression models such as linear regression with and without interactions, different regression trees, and support vector machine (SVM) regression models. GPR outperformed other models in training (both in terms of RMSE and *R*^2^; see Table 1).

**Table 1.** Comparison of different regression model performances.

| Model | RMSE | $R^2$ |
| --- | --- | --- |
| Linear Regression | 15831 | 0.85 |
| Decision Tree | 5616 | 0.98 |
| SVM | 6696 | 0.97 |
| GPR | 4990 | 0.99 |

### 3. Random Forest Details

Random Forest classifiers combine several decision trees that use multiple subsets of data from a training sample to produce better predictive performance than that of a single decision tree. The advantage of a Random Forest classifier is that it handles high dimensionality in data, as well as missing values. It works via the following principles: assume we have an observation *y*_*i*_ and the feature associated with it is *x*_*ij*_. Here, *i* = 1, …, *N, j* = 1, …, *M* and *N* and *M* are the number of observations and features, respectively. We first take a subset from *N* number of training data randomly with replacement. We then take a subset of *M* features randomly. We split the node iteratively by finding the feature associated with the best split. With this iteration, we grow the largest tree. We then repeat these steps to aggregate *n* number of trees to create our Random Forest. We generated 30 trees using a five-fold cross-validation and an independent test set to validate our model. Random Forest was used as a plug-in to the FANCY in order to predict the assay type of the data from the sequencing statistics in the case that metadata is missing.

### 4. Dataset

We used RNA-Seq data from 432 individuals generated by the gEUVADIS project [6], H3K27ac ChIP-Seq data from 100 individuals generated by the PsychENCODE Consortium [7] ATAC-Seq data from 344 individuals generated by the BrainGVEX project [7], and ATAC-Seq data from 288 individuals generated by the PsychENCODE Consortium [7]. We then used the GATK Best Practices from RNA-Seq and DNA data [8,9] to call SNVs and small insertions and deletions (Figure 3, Table 2). We treated each chromosome separately, which resulted in 25,152 data points. We randomly divided the data in half to use as training and test sets. We also separately validated our model by using 308 data points from the RNA-Seq study by Kilpinen et al. [10].

**Table 2.** Comparison of different regression model performances when the data is sub-sampled.

| Model | RMSE | $R^2$ |
| --- | --- | --- |
| Linear Regression | 11003 | 0.88 |
| Decision Tree | 7898 | 0.94 |
| SVM | 11722 | 0.86 |
| GPR | 5544.3 | 0.97 |

**Fig. 3.**
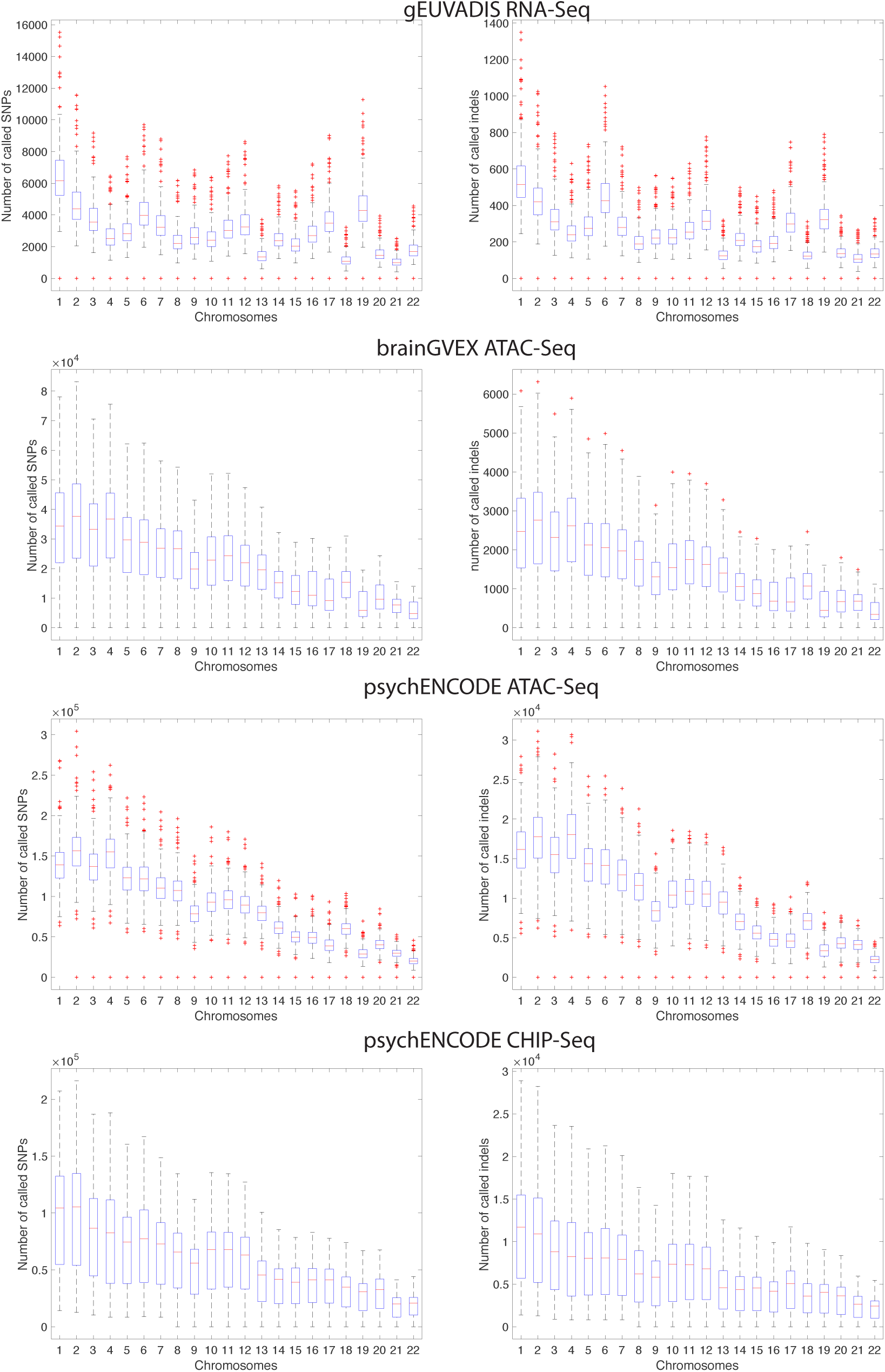
Distribution of the number of called SNPs and indels from functional genomics data. We used the GATK Best Practices from RNA-Seq and DNA data to call SNVs and small insertions and deletions.

In total, we had 13,537 data points from ATAC-Seq, 9,456 data points from RNA-Seq, and 2,159 from ChIP-Seq. In order to understand whether an imbalance in the number of categories affected our model selection, we repeatedly sub-sampled 2,159 data points from the RNA-Seq and ATAC-Seq categories and trained multiple regression models. We found that GPR is the best regression model in each case (Table 2).

## Results

Genotyping from DNA sequences is the process of comparing the DNA sequence of an individual to that of the reference human genome. To be able to successfully genotype, one needs a substantial depth of sequencing reads for each base pair. According to the Lander-Waterman statistics for DNA sequencing, when random chunks of DNA are sequenced repeatedly, the depth per base pair follows a Poisson distribution with a mean that can be estimated from the read length, number of reads, and the length of the genome [11]. For example, as RNA-Seq aims to sequence expressed genes, one would expect that sequencing depth per base pair does not follow Poisson statistics. Genotyping using reads from RNA-Seq experiments is biased towards variants that are in the exonic regions. Conversely, ChIP-Seq is biased against RNA-Seq, as it targets non-coding genome such as promoters and enhancers (see Figure 1c).

We hypothesized that the statistical properties of the depth per base pair distribution are strong indicators of the number of variants that can be inferred from functional genomics data. We used a total of six sequencing features: 1) the average depth per base pair 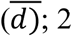 the total fraction of the genome that is represented at least by one read (i.e., the breadth, 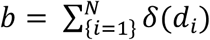, such that *δ*(*d*_*i*_) = 1 if *d*_*i*_ > 0, *b* = 0 otherwise and *N* is the total number of nucleotides in the genome); 3) the standard deviation of the depth distribution; 4) skewness (i.e., whether the distribution is larger on the right or left side of the mean); 5) kurtosis (i.e., whether or not the depth distribution has big tails); and 6) the type of the experiment (i.e., RNA-Seq, ATAC-Seq, or ChIP-Seq).

FANCY predicts the cumulative number of leaking SNVs with an *R*^2^ of 0.99 for training (with five-fold cross-validation) and 0.90 for independent test RNA-Seq, 0.99 for independent test Assay for Transposase-Accessible Chromatin using sequencing (ATAC-Seq), and 0.99 for independent test ChIP-Seq datasets (Figure 4a, Table 3). We used mean squared error as our loss function in the regression model [see Table 1-3 for Root Mean Squared Error (RMSE)]. Our predictions are in strong agreement with the true number of leaking variants in all the independent test datasets (Figure 4a). To easily interpret the performance of our predictions, we calculated the deviation from the true values by calculating (predicted value - true value) / true value. The negative values indicate underprediction (i.e. the number of predicted SNVs is lower than the true number of leaking SNVs). On average, we had 8% prediction error for all of the independent test sets (Figure 4b).

**Table 3.** The maximum and minimum number of variants leaked in each experiment and the RMSE of our predictions in these test datasets.

| Assay | number of test data points | max. number of variants | Min. number of variants | RMSE |
| --- | --- | --- | --- | --- |
| RNA-Seq | 4740 | 12,928 | 379 | 422.76 |
| ATAC-Seq | 6753 | 238,481 | 65 | 4503.10 |
| ChIP-Seq | 1082 | 210,657 | 2665 | 5381.22 |

**Fig. 4.**
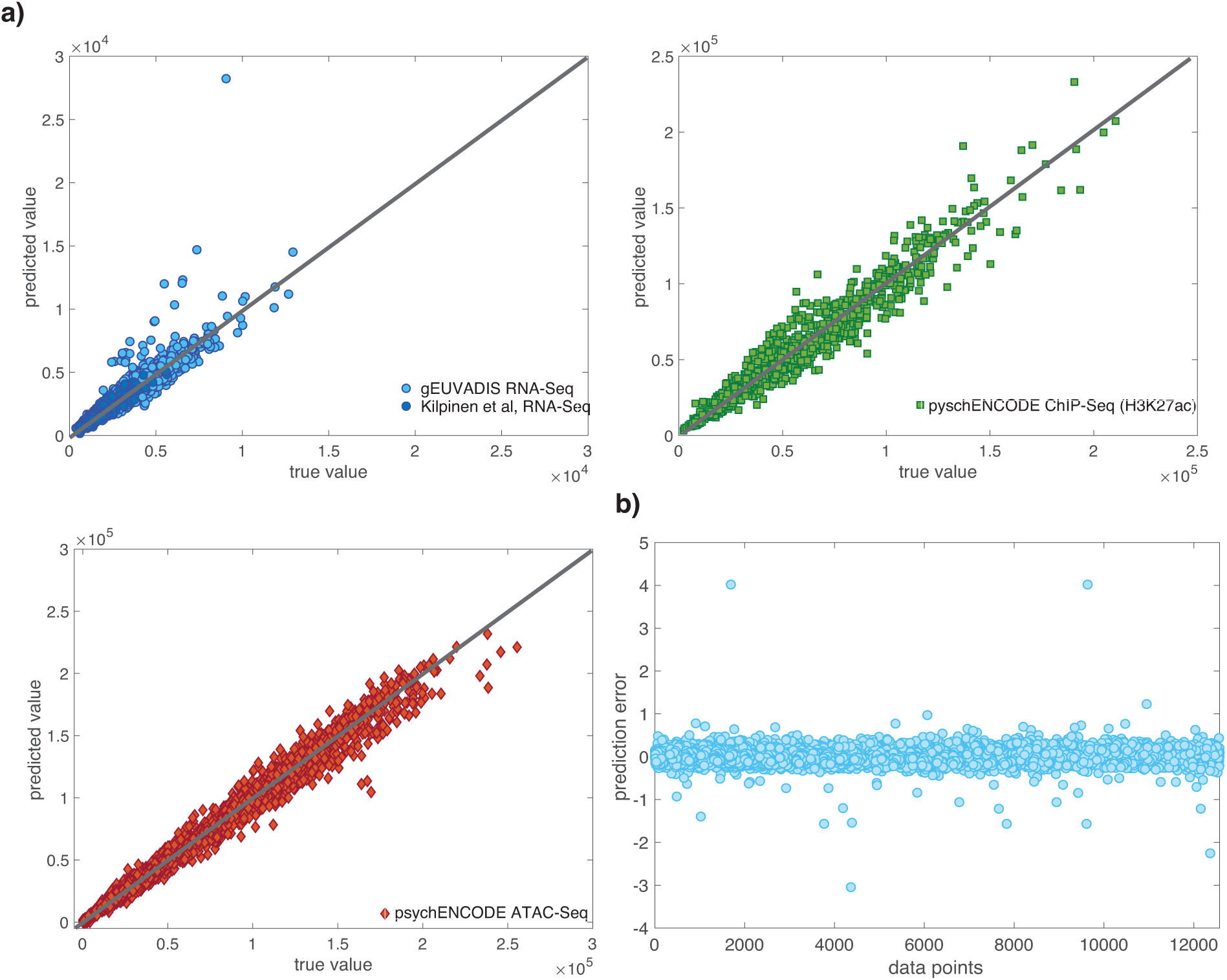
Details of FANCY. (a) The performance of FANCY on the independent test datasets. (b) The ratio of the error in the predictions with respect to true values. Negative values indicate that the predicted values are lower than the true values, and positive values indicate the predicted values are greater than the true values. Zero indicates perfect prediction.

If a functional genomics experiment is leaking more than 1,000 variants, the associated privacy risk for re-identification is at the maximum, regardless of the absolute value of the number of variants. Thus, the risk assessment is more valuable for experiments that are leaking a low number of variants, as mis-predicting these values only slightly may result in the release of private data. Thus, we developed another regression model (see Methods) that aims to predict the leakage more precisely when the number of leaking variants is low. This second model had an RMSE of 75.64, 74.8, and 74.0 for independent test RNA-Seq, ATAC-Seq, and ChIP-Seq datasets, respectively, in which the maximum number of leaking variants is 1,000 (Figure 5). We also calculated the number of under-predicted (predicted value is lower than true value) leaking variants and found that we have no under-predicted leaking variants when the total number of leaking variants is lower than 400, and only three under-predicted leaking variants when the total number of leaking variants is between 400 and 500. Moreover, both FANCY and FANCY_low_ can also output the number of leaking variants within 95% confidence interval (Figure 6).

**Fig. 5.**
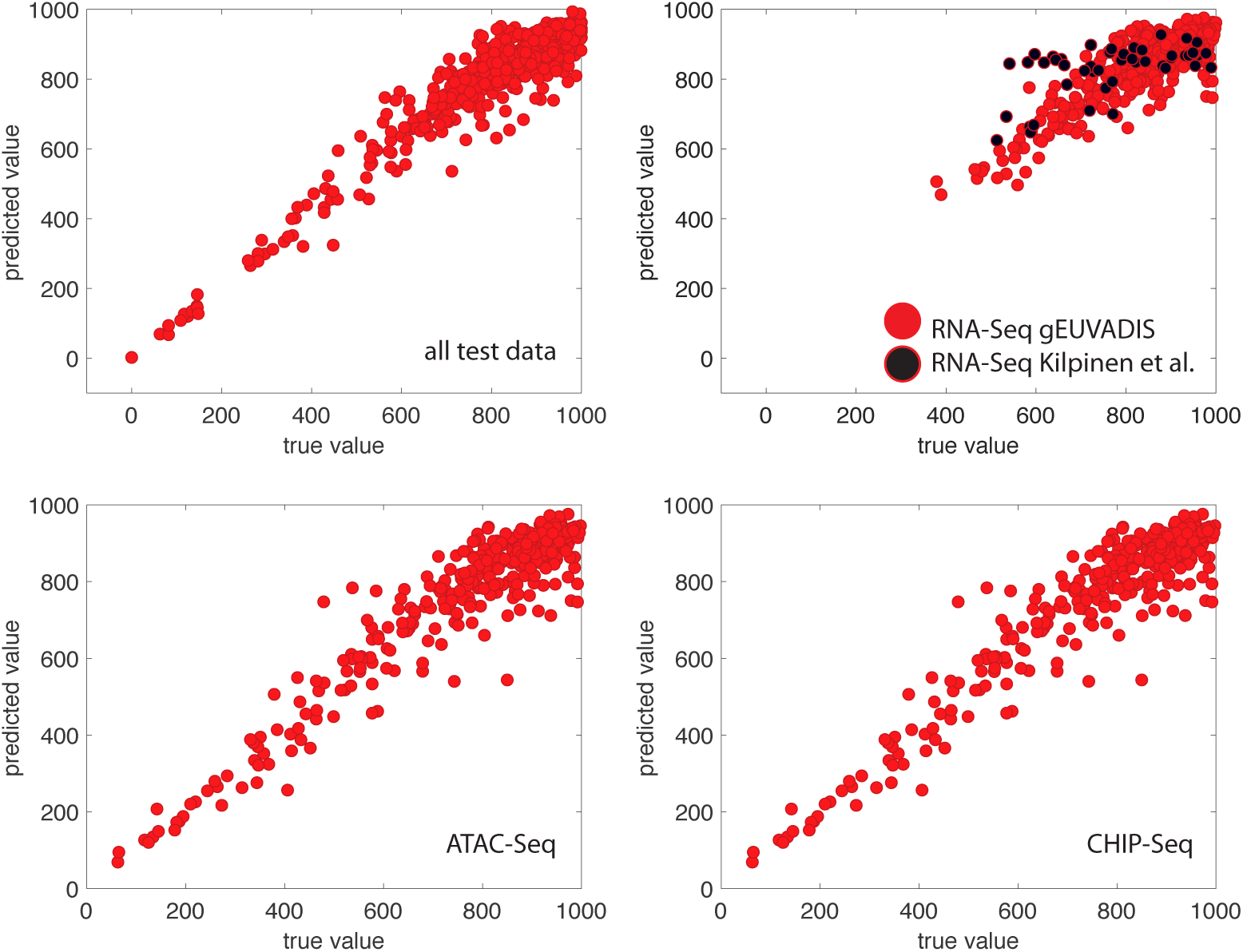
Performance of FANCY_low_ when the number of leaked variants is less than 1000. Performance for the test dataset, RNA-Seq, ATAC-Seq and ChIP-Seq are shown separately. For RNA-Seq, we validated our model with two datasets, shown in red and black.

**Fig. 6.**
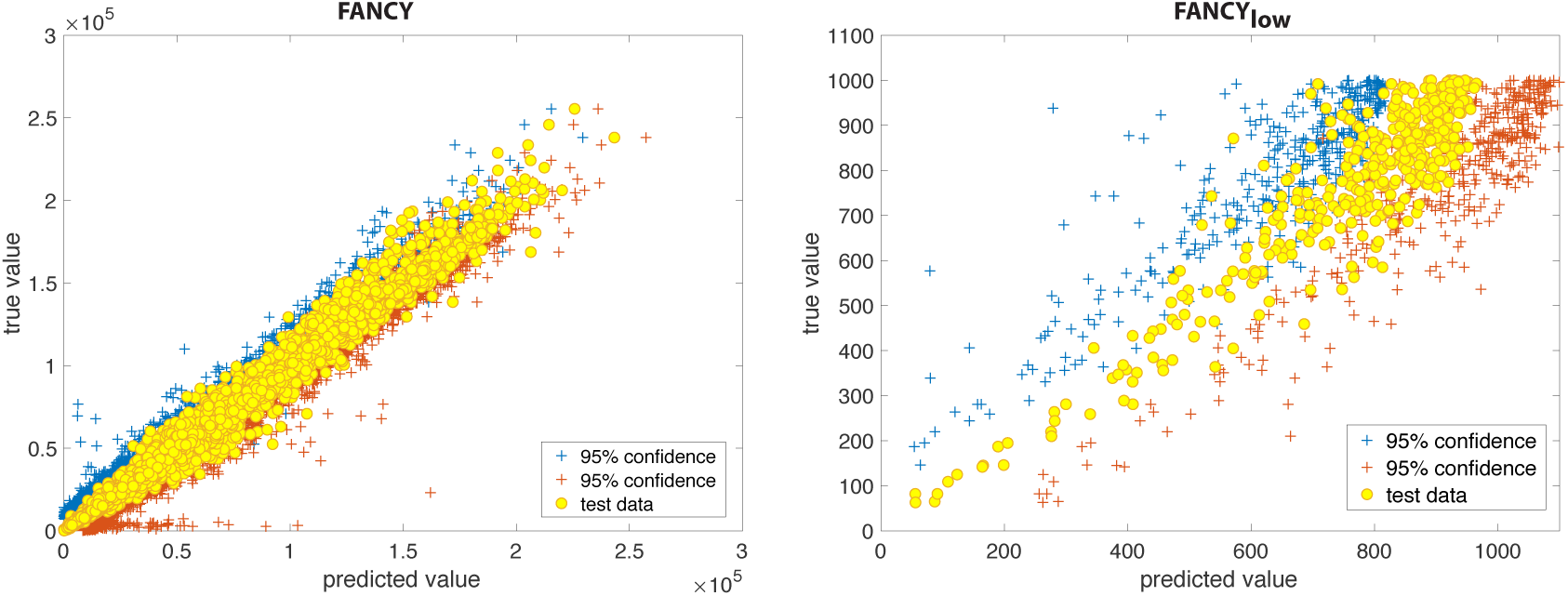
Predicted vs. true values within 95\% confidence level.

To further understand the role of the features on the prediction performance, we did a “leave one feature out” test and found that the mean depth 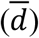 and breadth (*b*) of the coverage had the greatest effect on the performance of the predictor (Figure 7a). We then created predictors by using only (1) mean depth, (2) breadth, and (3) mean depth and breadth as the features. However, these predictors performed worse than the original model (Figure 7a). These results show that although breadth is the highest contributing feature, all of our features contributed to the final model; indeed, the RMSE is the lowest when we use all of the features (Figure 7a).

**Fig. 7.**
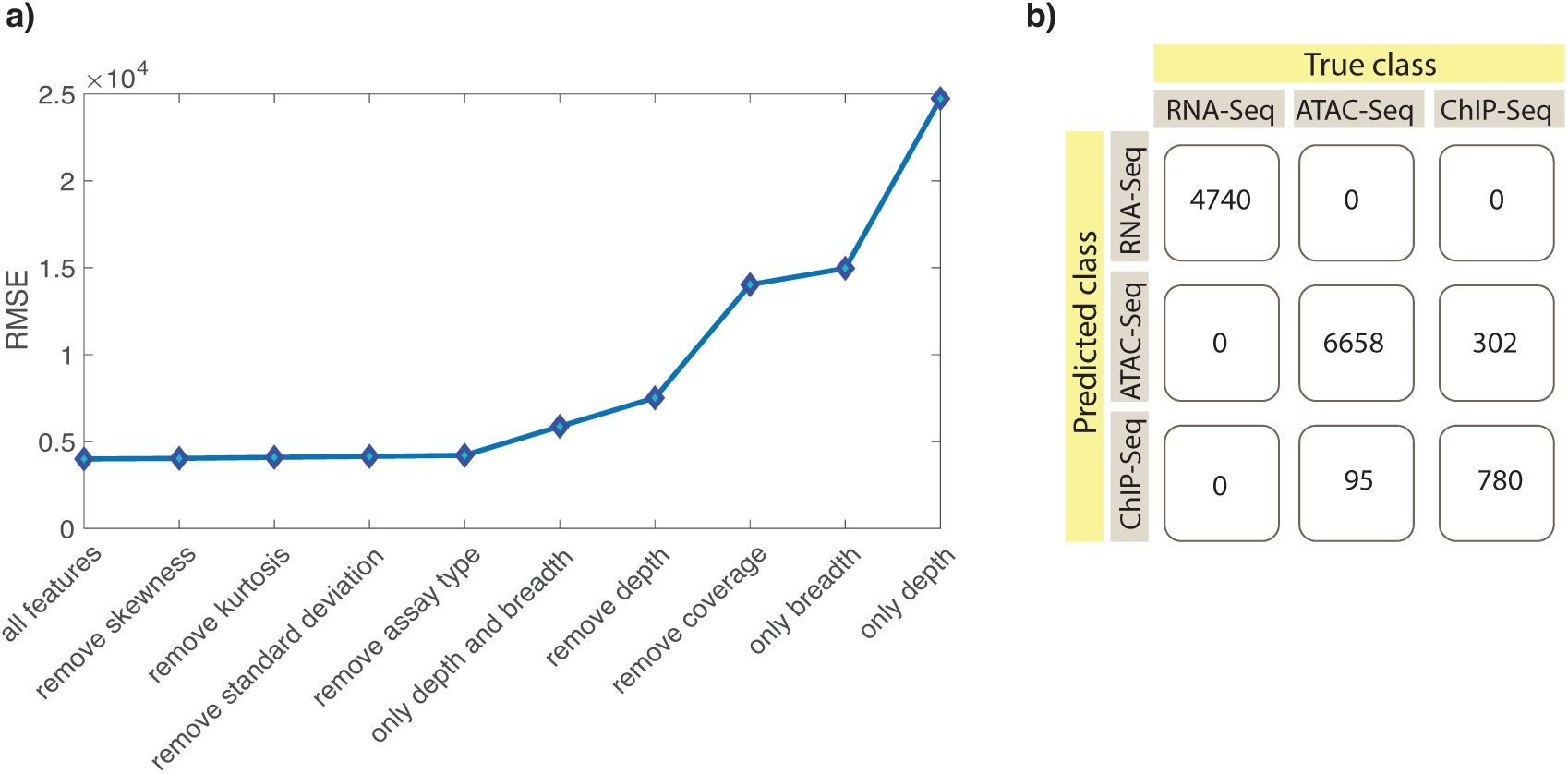
Feature importance and classifier performance. (a) Lower RMSE corresponds to better predictions. The difference between the first five points on the figure is smaller compared to other data points, however, lowest RMSE is still observed when using all the features in the model. (b) A Random Forest Classifier is developed to predict the type of the experimental assay by using sequencing statistics as features. Here we show the performance of this classifier.

In addition to estimating privacy risk, FANCY can also be used to plan functional genomics experiments (i.e., for a target number of SNVs, one can back-calculate the required sequencing statistics). Moreover, we also developed a Random Forest classifier as a plug-in that predicts the type of the assay (RNA-Seq vs. ATAC-Seq vs. ChIP-Seq) by using the sequencing statistics as features, which can be broadly useful to the community for characterizing samples with missing metadata. This classifier has an average accuracy of 96.8%, precision of 94.9%, recall of 90.2%, and F1 score of 93.3% (Figure 7b).

## Discussion

How can we quantify the privacy risk that accompanies collection and sharing of functional genomics data? In this study we addressed this question with FANCY, a model using Gaussian Process Regression followed by rare vs. common variant estimation to predict the number of total leaking variants in a functional genomics dataset. We showed that this prediction can be generated with high accuracy without relying on genotyping or a gold-standard genotype list, which requires significant resources to obtain and may not always be obtainable for every individual. We determined that the depth and breadth of sequencing coverage had the greatest influence on the predictor, and that using all of the features yielded the most accurate prediction. Not only does FANCY quantify the risk of privacy leaks from an existing dataset, but it also allows experimentalists to design functional genomics sequencing experiments, before collecting data, with the goal of obtaining a desired number of variants.

FANCY can also be used to accelerate sharing of functional genomics datasets. Functional genomics researchers may want to share their data alongside their published results. However, funding agencies’ data-sharing policies typically require extensive privacy risk assessments before the data is released. These assessments can be lengthy and costly, and can significantly delay data release (sometimes until after the relevant article has been published). Our tool will allow for fast assessments of privacy risk in functional genomics datasets, which will permit faster release of data. After FANCY, further assessments may be done of the data with significant risk.

If a functional genomics dataset leaks more than 1,000 variants, the risk of re-identifying an individual from that dataset is maximized. However, when the number of leaking variants is low, the qualitative measurements of sharing risk commonly given in consent documents may not be accurate and will not give the individual a sense of the true risk of privacy loss. Therefore, we designed FANCY_low_ a specially modified version of FANCY with improved precision just for a low number of leaking variants. Given that an individual’s privacy is at stake, it was important to minimize under-predictions of leaking variants as this could lead to a dataset being labeled safe to share when it actually permits re-identification.

Privacy protection is the core goal of developing FANCY and related tools, but they have other uses in genomics research. In cases where whole genome sequence data is missing, for example, FANCY can also be used to determine whether SNP calling is possible using a particular dataset. Additionally, our Random Forest Classifier, which we developed alongside FANCY, can be used to identify the kind of experiment a dataset came from in the case of missing metadata. This latter tool does not protect privacy, but helps to maximize data utility.

The privacy risk associated with human DNA sequencing has been acknowledged for years. Yet, as scientists have become increasingly interested in a more diverse set of experimental human omics data, it is critical to develop companion tools to assess the privacy risk of those data. In this study we contribute FANCY to the toolbox. Importantly, these tools must be convenient for scientists to use and easy for patients/individuals to understand. FANCY requires only a few files and can be run easily from the command line. Furthermore, we set up a simple web page, which allows users to input statistics about their dataset and runs FANCY to output a leakage prediction and qualitative privacy risk assessment. These user-friendly components are a key benefit of FANCY, and require no private information to be input to our servers, making it convenient for experimentalists to rapidly assess the risk of privacy before they release the data.

